# Evolutionary diversification of ancestral genes across vertebrates and insects

**DOI:** 10.1101/2024.06.11.598551

**Authors:** Federica Mantica, Manuel Irimia

## Abstract

**Background:** Vertebrates and insects diverged approximately 700 million years ago, and yet they retain a large core of conserved genes from their last common ancestor. These ancient genes present strong evolutionary constraints, which limit their overall sequence and expression divergence. However, these constraints can greatly vary across ancestral gene families and, in at least some cases, sequence and expression changes can have functional consequences. Importantly, overall patterns of sequence and expression divergence and their potential functional outcomes have never been explored in a genome-wide manner across large animal evolutionary distances.

**Results:** We focused on approximately 7,000 highly conserved genes shared between vertebrates and insects, and we investigated global patterns of molecular diversification driven by changes in sequence and gene expression. We identified molecular features generally linked to higher or lower diversification rates, together with gene groups with similar diversification profiles in both clades. Moreover, we discovered that specific sets of genes underwent differential diversification during vertebrate and insect evolution, potentially contributing to the emergence of unique phenotypes in each clade.

**Conclusions:** We generated a comprehensive resource of measures of sequence and expression divergence across vertebrates and insects, which revealed a continuous spectrum of evolutionary constraints among highly conserved genes. These constraints are normally consistent between these two clades and associated with specific molecular features, but in some cases we also identified cases of lineage-specific diversification likely linked to functional evolution.

## Background

Vertebrates and insects are two clades of bilaterian animals which diverged approximately 700 million years ago (MYA; [1]). On one hand, these clades exhibit relatively comparable body plans and homologous adult tissue types [2]. On the other hand, they also evolved unique biological traits [3–5], and are characterized by rather distinct genomic and molecular evolutionary rates [6,7].

Notwithstanding their large evolutionary distance, vertebrates and insects share a strong core of conserved genes, representing ∼25-65% of their protein-coding complement [8–10]. Since these genes are still recognizable as evolutionary related after 700 million years, their encoded protein sequence and expression profiles are likely subjected to strong evolutionary constraints. However, even within this set of highly conserved genes the degree of such constraints can greatly vary. Evolutionary constraints have a direct read-out in the overall levels of molecular diversification (in terms of sequence and/or expression divergence) within each given gene family, and are generally expected to be determined by their different functional requirements. For instance, genes encoding highly structured proteins usually have greater sequence constraints than those with abundant intrinsic disordered regions [11,12]. Similarly, genes with neural-specific expression in vertebrates show higher expression conservation than those specifically expressed in other tissues [10,13,14].

Remarkably, gene duplication can radically alter these intrinsic constraint patterns, both at the sequence and expression levels. Accordingly, highly duplicated genes generally exhibit greater molecular diversification compared to single-copy orthologs [10,15–17]. While this increased molecular diversification is normally due to the reduced constraints upon duplication, and it is thus expected to have a neutral effect, in at least some cases such changes lead to the evolution of new functional properties. Therefore, even if this functional interpretation applies only to a limited number of events, understanding patterns of molecular divergence in a gene family may guide sound hypotheses about its functional evolution.

Notably, while changes in sequence or expression alone can drive functional evolution, synergistic modifications at both levels may be more efficient to this aim. Indeed, the combined effect of sequence and expression variation in determining functional evolution has been proved by single-species studies for several groups of paralogs [18–21]. However, sequence and expression changes within gene families have never been investigated together across large animal phylogenies and in a genome-wide manner. On one hand, some landmark studies reconstructed global patterns of gene gains, losses and duplications throughout animal evolution [8,9,22], but they neither investigated sequence divergence within gene families, nor complemented their findings with expression data. On the other hand, large comparative transcriptomic studies normally focus on sets of conserved (often 1:1) orthologs [10,13,14,23,24], but do not usually integrate evolutionary expression changes with information about sequence evolution.

In this work, we use novel measures of sequence and expression divergence to investigate global patterns of molecular diversification of ancestral gene orthogroups within and between vertebrates and insects, with the assumption that they could be informative not only about their molecular constraints but also regarding trends of functional evolution in the two clades. We focused on a set of ∼7000 highly conserved gene orthogroups, characterizing general features associated with molecular diversification and identifying groups of genes that exhibit common or unique diversification patterns across vertebrates and insects. Finally, we determined potential connections between molecular diversification and tissue-related attributes, highlighting the important role of highly conserved genes for the evolution of clade-specific phenotypic traits.

## Results

### Global rates of sequence and expression diversification in vertebrates and insects

We selected a symmetric phylogeny of eight gnathostome vertebrates and eight insects, with pairs of vertebrate and insect species located at equivalent phylogenetic positions on the two main branches (**Fig. 1a**). As it included species with similar divergence times, our phylogenetic tree was optimal to directly compare the evolution of molecular traits between the two clades. We specifically focused on protein-coding genes broadly conserved across this phylogeny (6,787 orthogroups; [10]). For all these genes, we extracted (i) their encoded protein sequence and (ii) their expression profile across seven homologous tissue types (i.e., neural, testis, ovary, muscle, excretory system, epidermis and guts) (see **Methods**). We then used these measures to compute, respectively, the sequence and expression similarities of homologous genes either within vertebrates or within insects (**Fig. 1b,c, Supplementary Fig. 1** and **Supplementary Table 1;** see **Methods**), which we used as proxies of molecular diversification in the corresponding gene orthogroup.

The overall distributions of sequence similarities revealed that protein sequences are in general more conserved in vertebrates compared to insects (**Fig. 1d**; two-sided Wilcoxon’s test, p-value < 2e-16). This is partially expected because of the shorter generation times [7] and the bigger population sizes [25,26] of insect species compared to vertebrates. On the contrary, expression profiles across tissues show similar conservation levels between the two clades (**Fig. 1e**). This is in line with previous studies, which demonstrated how transcriptional networks evolve with largely comparable rates between mammals, birds and insects [27], or how expression divergence reaches a relatively quick plateau with increasing evolutionary distances, both within mammals [13,28,29] and fruit flies [30]. In summary, while sequence changes seem to have been overall more prevalent in insects compared to vertebrates, expression changes contributed similarly to the modification of the ancestral molecular landscape in the two clades.

**Fig. 1:**
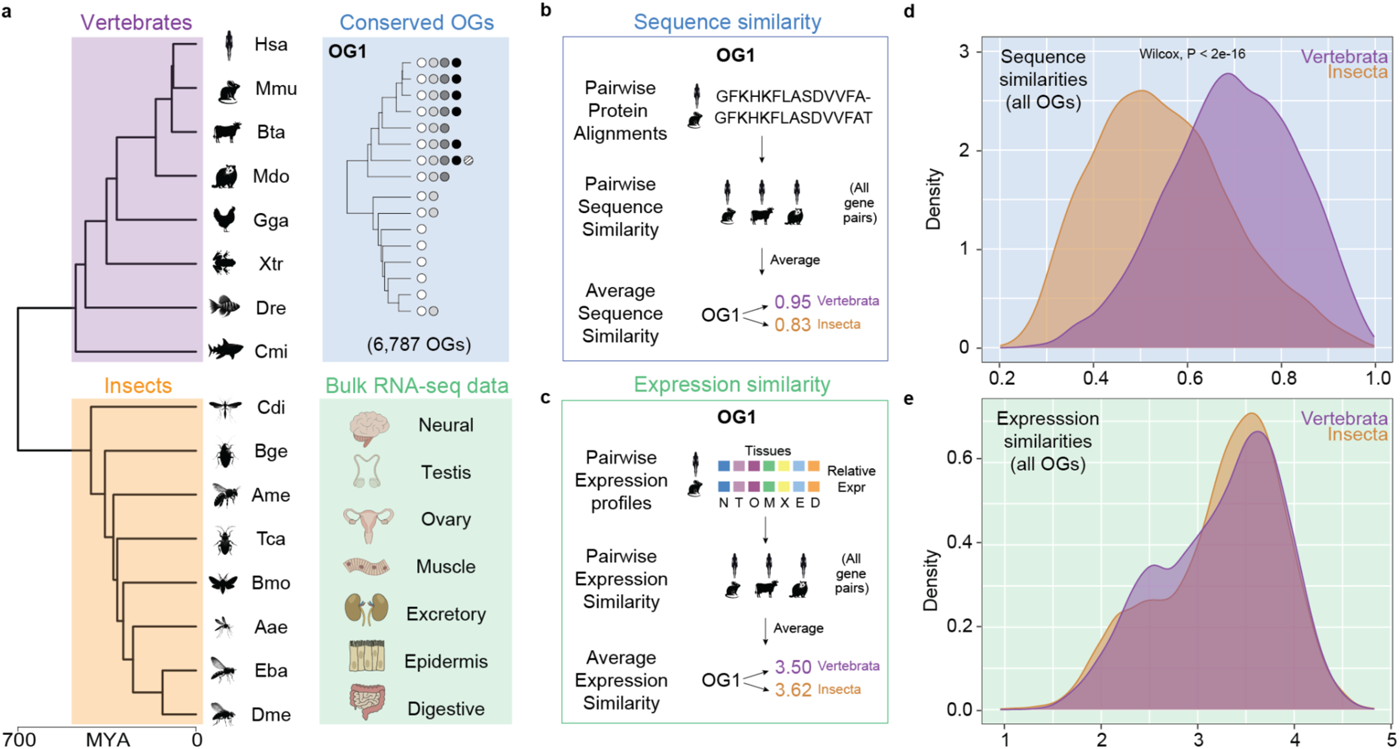
Framework overview and definition of sequence and expression similarities: **a.** Left: phylogenetic tree including the scientific acronyms of the 8 vertebrate and 8 insect species considered in this study. *Hsa*: human, *Mmu*: mouse, *Bta*: cow, *Mdo*: opossum, *Gga*: chicken, *Xtr*: tropical clawed frog, *Dre*: zebrafish, *Cmi*: elephant shark. *Dme*: fruit fly, *Eba*: marmalade hoverfly, *Aae*: yellow fever mosquito, *Bmo*: domestic silk moth, *Tca*: red flour beetle, *Ame*: honey bee, *Bge*: cockroach, *Cdi*: mayfly. (see **Methods** for corresponding scientific names). Top right: example of one the 6,787 considered gene orthogroups. Bottom right: tissues represented in our bulk RNA-seq dataset. Evolutionary distances were derived from timetree [1] (MYA: million years ago) and animal silhouettes were generated through *Bing Chat* by Microsoft (2023) https://www.bing.com/search. **b, c.** Scheme for the computation of the sequence (b) and expression (c) similarity measures. The procedure was performed separately for vertebrates and insects, returning two values per orthogroup. See **Supplementary** Fig. 1a for step-by-step schematics. **d, e.** Distributions of the sequence (d) and expression (e) similarity values for all gene orthogroups (n=6,787) within vertebrates (purple) and insects (orange).

### Comparisons of sequence and expression diversification within and between clades

Next, we separately compared rates of sequence or expression evolution of individual gene orthogroups between the two clades. In general, gene orthogroups show significant and positive correlation both in terms of sequence similarities (Person’s correlation coefficient: 0.631, p-value < 2e-16; **Fig. 2a**) and expression similarities (Pearson’s correlation coefficient: 0.559, p-value < 2e-16; **Fig. 2b**). This suggests that genes that tend to be more or less conserved in vertebrates also evolve with similar relative rates in insects, both from the sequence and the expression perspectives. Thus, the disposition to undergo molecular diversification driven by either type of molecular change seems to be, to a large extent, an intrinsic potential of each gene orthogroup, which is independently but comparably fulfilled in the two clades.

Moreover, we found that conservation levels of these two molecular traits tend to be associated, as we detected positive and significant correlations between sequence and expression similarities both within vertebrates (Pearson’s correlation: 0.401, p-value < 2e-16; **Fig. 2c**) and within insects (Pearson’s correlation: 0.377, p-value < 2e-16; **Fig. 2d**), in line with what was previously shown for more closely related species [28,31,32]. However, it should be noted that this association between sequence and expression conservation did not emerge from other studies [33,34], potentially due to the caveats of using correlations as measures of expression divergence [35] (see **Supplementary Discussion** for a comparison between alternative metrics of sequence and expression conservation). Importantly, despite their positive and significant association, sequence and expression similarities show a substantial deviation from perfect alignment (**Fig. 2c,d**). This indicates that, although genes predisposed to molecular diversification experience generally increased levels of both sequence and expression alterations, the exact interplay between these factors may vary across gene orthogroups and between clades.

**Fig. 2:**
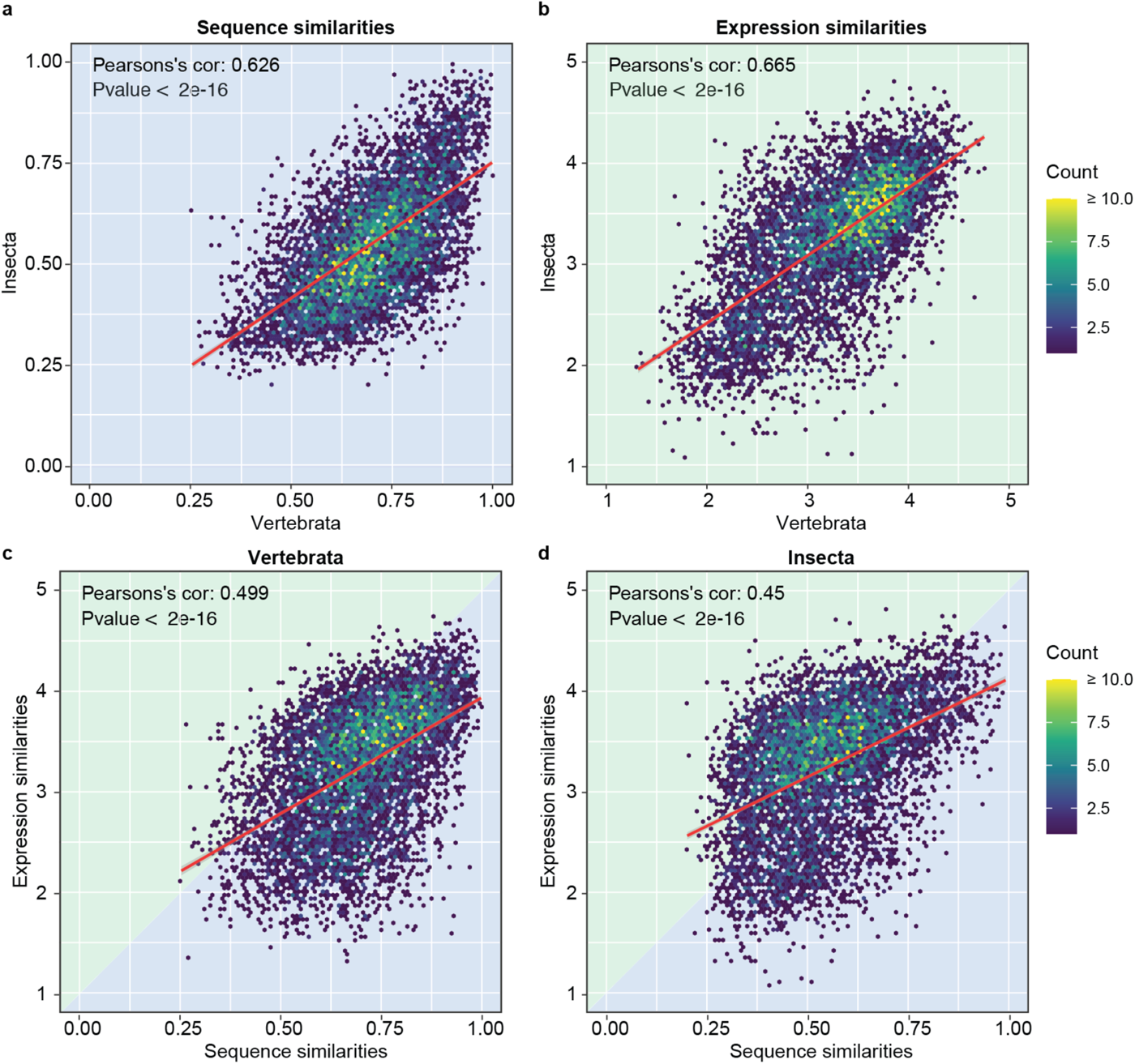
Correlation of sequence and expression similarities between and within clades. **a,b.** Correlation of the sequence (a) and expression (b) similarities of all gene orthogroups between vertebrates (x axis) and insects (y axis) (n=6,787). **c, d.** Correlation between the sequence (x axis) and expression (y axis) similarities of all gene orthogroups within vertebrates (c) and insects (d) (n=6,787).

### Features associated with molecular diversification

We then used sequence and expression similarities to define groups of highly or lowly diversified genes within vertebrates and within insects. In particular, we selected the 500 gene orthogroups with the lowest/highest sequence or expression similarities in each clade, defining eight groups of genes with the most extreme diversification profiles (**Fig. 3a** and **Supplementary Table 2**). According to our assumptions, highly diversified gene orthogroups should include the ones with weaker overall constraints, but also those that have more often undergone functional evolution; on the other hand, lowly diversified orthogroups are highly constrained and more likely to have strictly preserved their ancestral functions.

First, in-depth characterization of these orthogroups revealed that highly and lowly diversified genes were associated with significantly less and more lethal phenotypes, respectively, compared to all genes (one-sided Fisher’s tests: all p-values ≤ 0.001, except for the non-significant p-value of the vertebrate highly diversified orthogroups in terms of sequence; **Fig. 3b**). Second, and in part consistent with that observation, they also showed significantly higher and lower duplication levels compared to the background (one-sided Fisher’s tests: all p-values ≤ 0.0005; **Fig. 3c**). This pattern was particularly strong for vertebrates, most likely determined by the two rounds of whole genome duplications that characterize their early evolutionary history [36]. Moreover, these increased duplication levels were observed for highly diversified genes derived from both sequence and expression-based classifications, in line with the idea that gene duplication releases the evolutionary constraints of ancestral genes and potentially triggers their functional evolution through the combination of different molecular mechanisms [37].

Finally, we used a measure of tissue-specific expression known as Tau [38] to test for potential biases in tissue-specificity among the selected genes. All highly and lowly diversified gene groups were, respectively, significantly more and less tissue-specific than all genes together (one-sided Wilcoxon’s test, all p-values < 2e-16; **Fig. 3d**). On one hand, this was consistent with the concept that orthogroups with minimal mutation load will include mainly essential housekeeping genes, characterized by broad expression profiles [39]. On the other hand, it also indicated that gains of tissue-specificity might be among the expression changes most employed to generate molecular diversification, and that a substantial proportion of this diversity might contribute to the evolution of tissue-related traits (in line with [10,13,14,28,29,40,41]). Importantly, the fact that also the highly diversified genes from the sequence-based classification show a dramatic increase in tissue-specificity is another strong indicator that both sequence and expression modifications might cooperate in determining functional evolution.

**Fig. 3:**
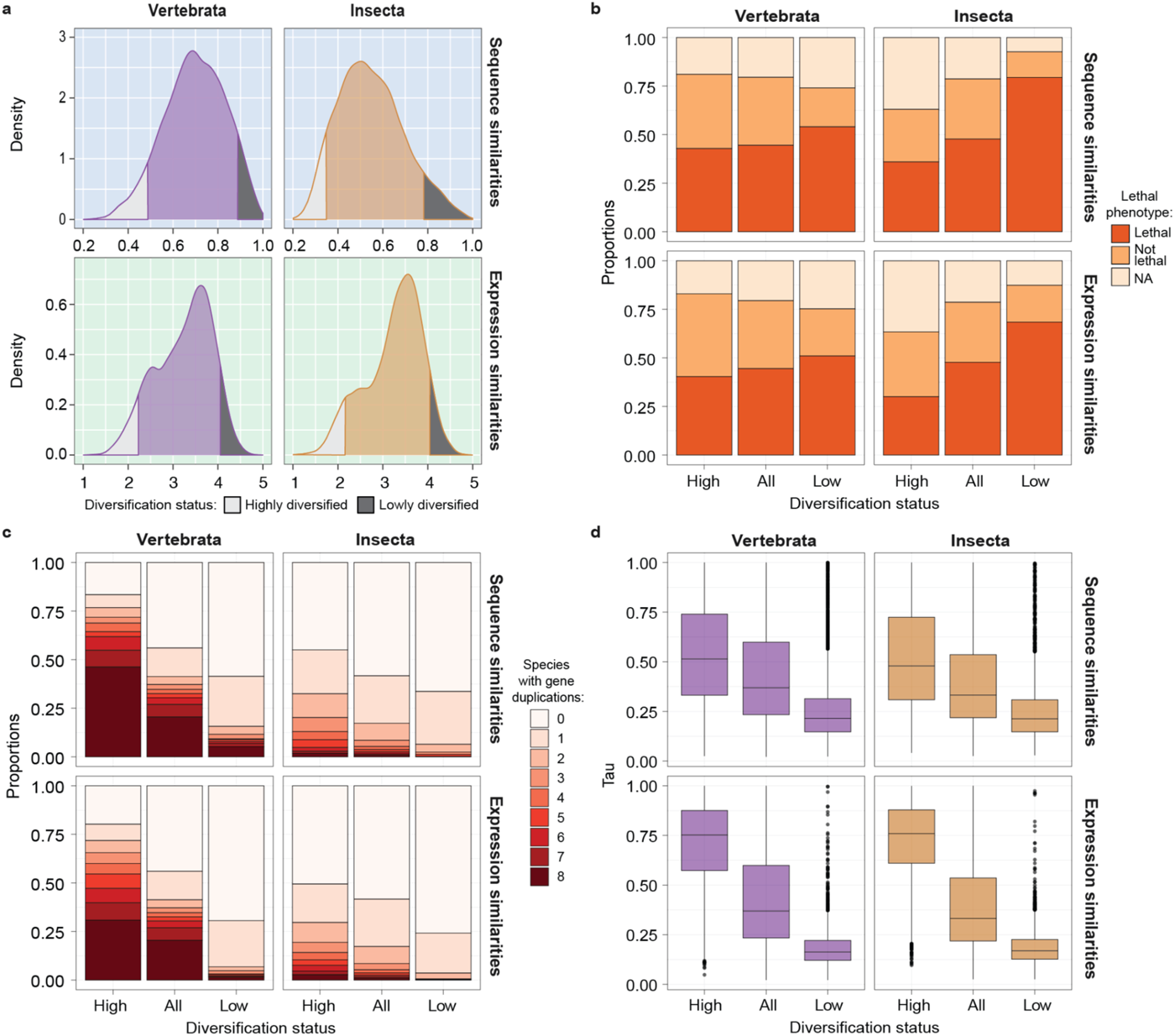
Characterization of highly and lowly diversified gene orthogroups. **a.** Distribution of the sequence (top) and expression (bottom) similarity values of all gene orthogroups in vertebrates (left; purple) or insect (right; orange). The highly (n=500) and lowly (n=500) diversified gene orthogroups are highlighted in white and black, respectively. **b.** Proportions of highly diversified, all and lowly diversified orthogroups associated with a lethal, non-lethal and uncharacterized phenotype. Each plot refers to the corresponding distributions in panel a. See **Methods** for phenotype definition. **c.** Duplication profiles of highly diversified, all, and lowly diversified orthogroups, where species with duplications are defined as all species with at least two conserved paralogs in a given orthogroup. Each plot refers to the corresponding distributions in panel a. **d.** Distribution of Tau values for genes included in highly diversified, all and lowly diversified orthogroups. Each plot refers to the corresponding distributions in panel a.

### Functional categories with common diversification patterns between clades

Then, we focused on those genes that undergo similar extreme levels of molecular diversification (very low or very high) in both vertebrates and insects and through the combined action of sequence and expression changes. First, we selected all the genes belonging to at least three out of four groups of lowly diversified genes (core of the Venn diagram, **Fig. 4a** and **Supplementary Table 3**), which should include genes that mainly preserve highly ancestral functions. As expected, a GO enrichment analysis on these genes returned categories related to basic and highly conserved cellular organelles (e.g., nucleus, ribosome, nucleolus), molecular functions (e.g., rRNA binding) or biological processes (e.g., translation, mRNA/rRNA processing and splicing) (**Fig. 4b** and **Supplementary Table 2**).

Second, we repeated the same analysis for gene orthogroups belonging to at least three out of four groups of highly diversified genes (core of the Venn diagram, **Fig. 4c** and **Supplementary Table 3**), representing orthogroups with pervasive patterns of molecular diversification. These genes showed clear enrichments in various cilium-related categories (e.g., axoneme), gamete generation and extracellular regions (**Fig. 4d** and **Supplementary Table 3**). Many of these enrichments presumably arise from the great variation of male reproductive systems in animals, which are the result of a strong selective sexual and molecular pressure [24,42]. For example, the axoneme enrichment likely reflects how this ancient microtubule-based structure, a core component of spermatozoa flagella in most animal species [43], was diversified across a wide range of reproductive niches.

**Fig. 4:**
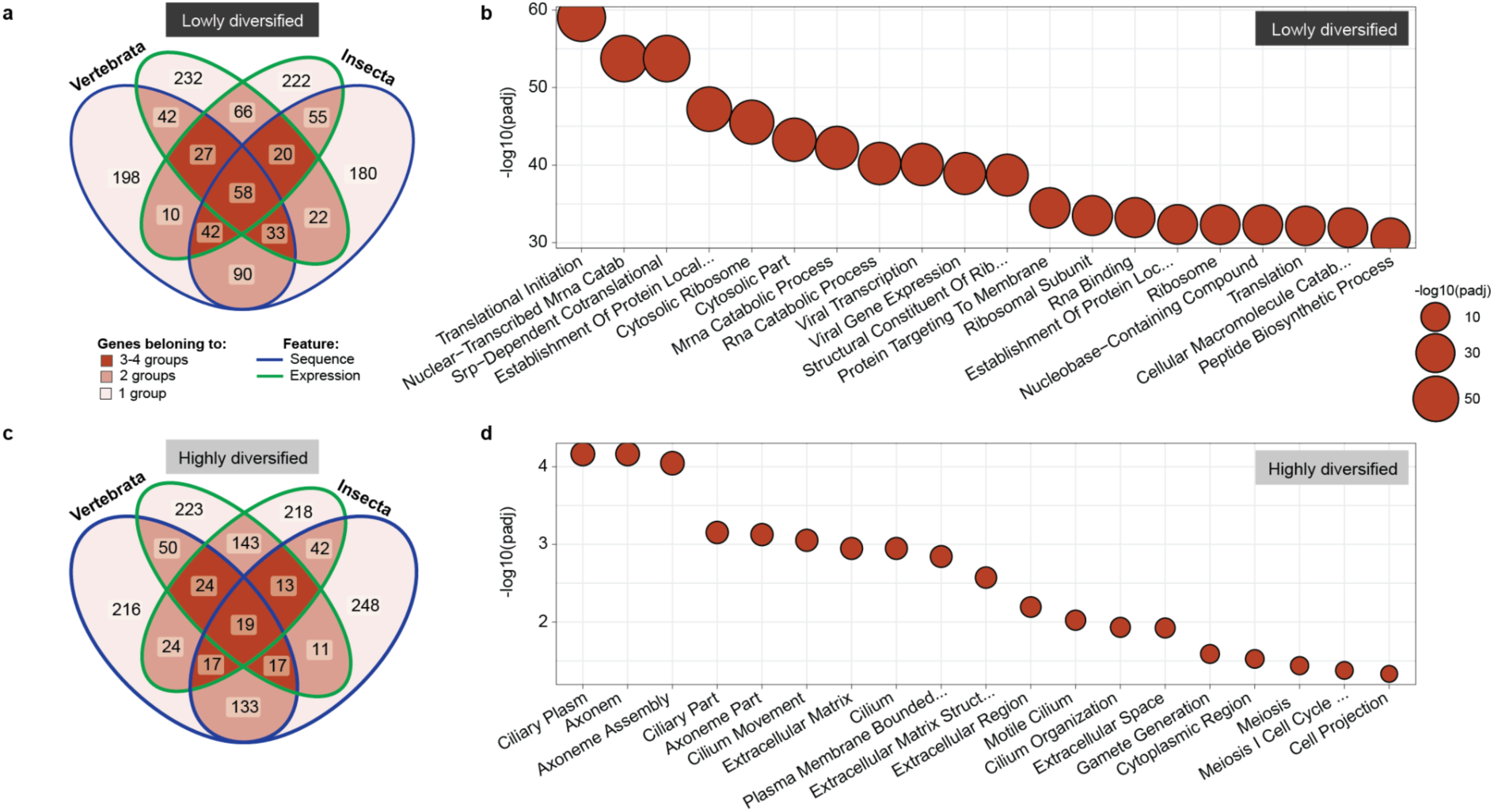
Functional categories with common diversification patterns between clades. **a,c.** Venn diagrams representing the overlap between lowly (a) and highly (c) diversified gene orthogroups defined based on vertebrate/insect sequence (blue) or expression (green) similarity. A common legend for both panels is depicted in panel a. **b,d.** Significant categories (FDR corrected p-values ≤ 0.05, intersection ≥ 5, precision ≥ 0.05) from a GO enrichment analysis performed on the lowly (b) and highly (d) diversified gene orthogroups common to 3-4 groups depicted in panel a and c, respectively (center of the Venn diagrams). Results are limited to the top 20 categories, but full reports are available in **Supplementary Table 3.**

### Functional categories with differential diversification patterns between clades

We next focused on gene orthogroups that are more divergent in one clade compared to the other, indicating that they have been differentially impacted by sequence or expression changes during vertebrate and insect evolution. In order to characterize these orthogroups, we computed the deltas of the respective sequence or expression similarities between the two clades (vertebrates minus insects; **Supplementary Fig. 2a,b** and **Supplementary Table 1**), which showed moderate but significant correlation across all orthogroups (Pearson’s correlation coefficient: 0.349, p-value < 2e-16, **Supplementary Fig. 2c**). Given the non-overlapping initial distributions of sequence similarities between vertebrates and insects (**Fig. 1d** and **Supplementary Fig. 2a**), we applied a z-score transformation to the deltas distribution of sequence and, for coherence, of expression similarities **(Supplementary Fig. 2d,e)**. In this way, the sign of the z-scored deltas is reflective of higher diversification rates in insects (positive values) or in vertebrates (negative values), but preserve the same characteristics of the original distributions (**Supplementary Fig. 2d-f**). We then performed a separate gene set enrichment analysis (GSEA) on these z-scored deltas for the sequence and expression similarities (**Fig. 5a** and **Supplementary Table 3**), highlighting all genes belonging to significant categories (FDR-corrected p-value ≤ 0.01) in **Fig. 5b,c**.

In general, genes with a diversification bias in insects were enriched for several cilium and cytoskeleton-related GO categories, especially when this bias was expression-driven. In contrast, genes with a diversification bias in vertebrates were enriched for functions associated with synaptic structures, ion transport and actin cytoskeleton. Importantly, while both vertebrate and insect genes with diversification biases tend to be more tissue-specific than their orthologs in the other clade (**Fig. 5d**), they most likely manifest their potential for functional evolution in different tissue contexts (**Fig. 5e**). On one hand, the insect group is mainly enriched in testis-specific genes, possibly reflecting the highly variable and specialized ciliary structure of their reproductive traits [44,45], and consistent with the GSEA results. On the other hand, neural-specific genes are strongly over-represented in the vertebrate group, also in line with their observed enrichment for neuronal functional categories and the overall increased complexity of the vertebrate brain [46,47]. All together, this analysis suggests that at least part of this molecular diversification is likely to result in functional evolution, and might have been differentially used in vertebrates and insects to shape some of their distinct tissue-related traits.

**Fig. 5:**
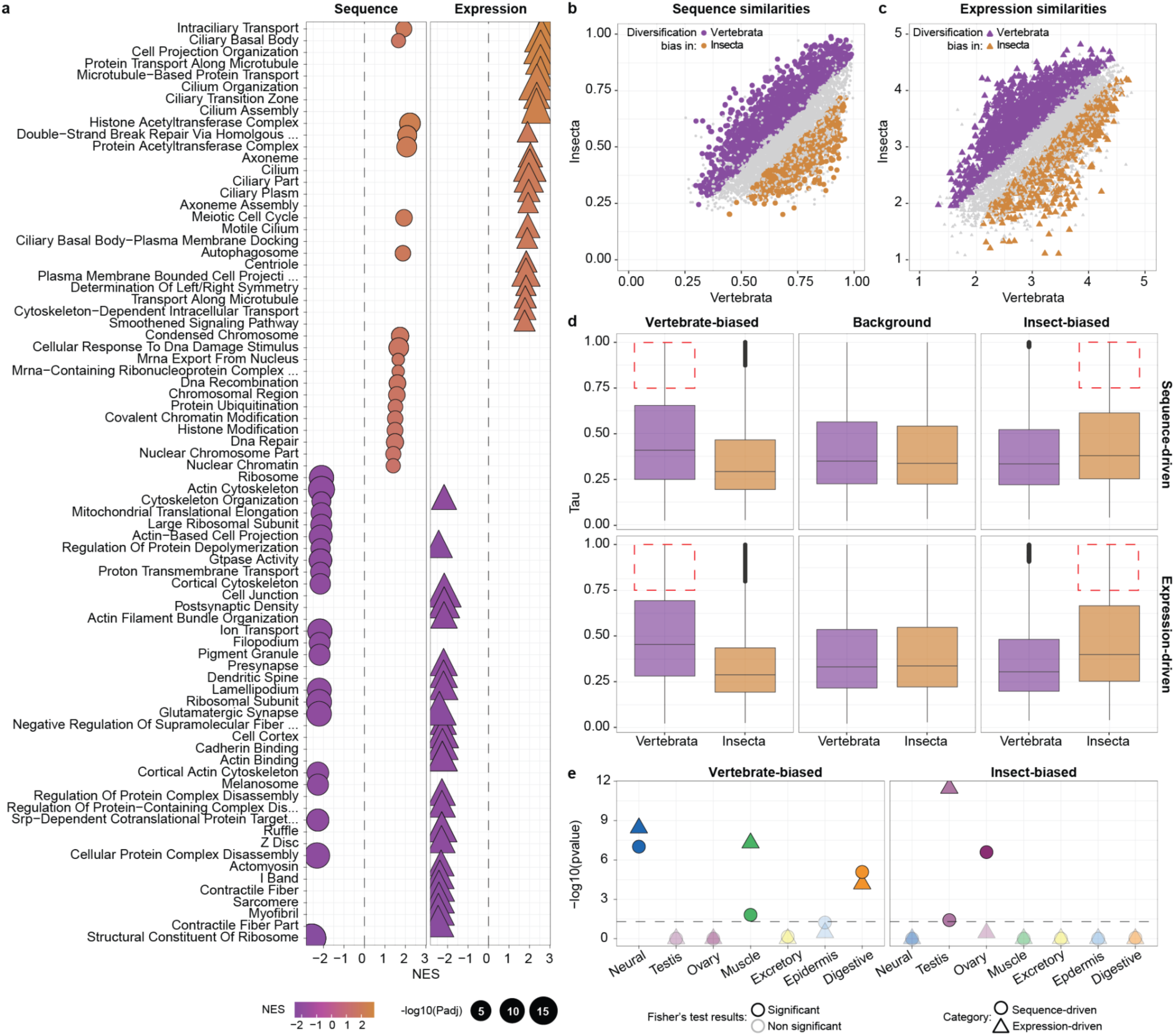
Functional categories with diversification biases between clades. **a.** Top significant categories (FDR-corrected p-values ≤ 0.01) from a GSEA performed on the z-scored deltas of sequence (left) or expression (right) similarities between vertebrates and insects. **b,c**. Same plots as in Fig. 2a**,b** but where the genes driving all significant enrichments are highlighted. The genes driving positive enrichment scores (high deltas, more diversified in insects) are highlighted in orange, while those driving negative enrichments (low deltas, more diversified in vertebrates) are highlighted in purple. **d.** Distribution of Tau values for genes in the orthogroups with diversification biases highlighted in (b) (top row) and (c) (bottom row) and the relative background composed by all orthogroups. The Tau values are separately plotted for the vertebrate and insect genes included in each orthogroup. The dashed, red lines contain the genes used as input for the Fisher’s exact test depicted in (e). **e.** Results of Fisher’s exact tests from the *fisher.test* function in R (*alternative=”greater”*) comparing the proportions of tissue-specific genes with diversification biases (Tau ≥ 0.75, dashed lines in (d) and (e)) in each tissue to the same proportions in the background. Tests with p-values ≤ 0.05 are considered as significant. **Abbreviations**: NES, normalized enrichment score.

## Discussion

This work revolves around a core of ∼7,000 genes shared between vertebrates and insects, inherited from their last common ancestor approximately 700 MYA. Given their ancient origin, these genes are likely subjected to strong evolutionary constraints, which limit their divergence in terms of encoded protein sequence and expression profiles. However, even if all ancestral genes are somewhat constrained, they are expected to show some variability in their tolerance to mutations. In fact, some genes keep performing their function through a wide array of sequence or expression alterations, while others are extremely sensitive to modifications of their ancestral molecular state. Moreover, although the majority of fixed molecular alterations in these genes are likely neutral, we know that they can sometimes mediate functional diversification. Thus, characterizing genome-wide landscapes of evolutionary constraints of ancestral animal genes would not only reveal which genes are more or less tolerant to molecular alterations, but also potentially identify those cases where these alterations lead to functional evolution.

Methodologically, we developed novel measures of sequence and expression divergence that align with more standard metrics but offer a few advantages (see **Supplementary Discussion**). Moreover, they are directly comparable between clades thanks to the symmetric structure of our phylogeny, where the equal number of species with matching phylogenetic distances in the vertebrate and insect lineages remove eventual biases due to differential species sampling or evolutionary times. These measures revealed that (i) sequence and expression divergence are highly variable among gene orthogroups but (ii) homologous genes generally share similar evolutionary constraints across clades and molecular layers. Thus, the disposition to undergo molecular diversification driven by either type of molecular change seems to be an intrinsic potential of each gene orthogroup, that is independently but comparably fulfilled in vertebrates and insects.

Using this characterization of molecular diversification at the gene orthogroup level, we tried to determine what might drive differences in evolutionary constraints among these highly conserved gene families. While pinpointing causative factors is challenging, we identified some associated biological features. Genes with higher molecular diversification levels tend to belong to orthogroups with more duplicates, be more tissue-specific and less lethal compared to other genes. This supports the model where duplication lifts at least some of the constraints of the ancestral gene, making it less susceptible to lethal mutations. However, the increased tissue-specificity of more diversified genes suggests that at least part of this diversification might not simply be due to lack of constraints, but might have been positively selected because of its functional effect in specific tissue contexts.

When is molecular diversification more probably linked to a functional outcome? Given the overall high correlation in evolutionary constraints between vertebrates and insects, it could be argued that genes for which diversification levels significantly differ between the two clades have more likely undergone functional changes in one of those clades. Functional evolution of ancestral genes, particularly after gene duplication, can occur through different modalities: for instance, neofunctionalization indicates emergence of a completely novel function, while specialization implies the evolution of a slightly modified (or specialized) version of the ancestral role. Both novel and specialized functions are expected to be optimized for more restricted biological contexts (e.g., a particular tissue type) compared to more general ancestral functions, in line with the most common regulatory fates observed upon whole genome duplication [16,48–50]. Consistently, we found that gene orthogroups with a diversification bias in vertebrates were significantly enriched for neural-specific genes, whereas genes with higher molecular diversification levels in insects showed over-representation of testis-specificity. This contrast emphasizes that different tissues in vertebrates and insects are presumably most influenced by functional evolution of ancestral genes, and this process likely contributed to the unique complexity of vertebrate brains and to the highly variable features of insect reproductive systems.

## Methods

### Phylogenetic tree

The phylogenetic tree used in this publication is a subset of the phylogeny of bilaterian animals from [10], which we here restricted to the (eight) vertebrates and (eight) insects (sixteen species in total). The vertebrate and insect branches in the resulting phylogeny are monophyletic and symmetric, meaning that each vertebrate is paired with an insect located at an equivalent phylogenetic position and with relatively comparable divergence times. The vertebrate species include human (*Homo sapiens, Hsa*), mouse (*Mus musculus, Mmu*), cow (*Bos taurus, Bta*), opossum (*Monodelphis domestica, Mdo*), chicken (*Gallus gallus, Gga*), tropical clawed frog (*Xenopus tropicalis, Xtr*), zebrafish (*Danio rerio, Dre*) and elephant shark (*Callorhinchus milii, Cmi*). The insect species include fruit fly (*Drosophila melanogaster, Dme*), marmalade hoverfly (*Episyrphus balteatus*, *Eba*), yellow fever mosquito (*Aedes aegypti, Aae*), domestic silk moth (*Bombyx mori, Bmo*), red flour beetle (*Tribolium castaneum, Tca*), honey bee (*Apis mellifera, Ame*), cockroach (*Blattella germanica*, *Bge*), mayfly (*Cloeon dipterum*, *Cdi*). See [10] for details on the assembly and gene annotation versions for each of the species.

### Gene orthogroups

We considered the ancestral gene orthogroups defined by [10], which originally included 7,178 orthogroups widely conserved across bilaterian animals, and filter this set to select only the orthogroups that were conserved in at least two vertebrates and two insects (6,787 out of 7,178, 95% of the original orthogroups). For all the main analyses, we considered all paralogs from all species within each gene orthogroup, which allowed us to capture their overall diversification patterns. However, we also generated four distinct 1:1 orthogroup sets, each composed of a representative paralog per species (when present) selected according to different conservation criteria (**Supplementary** Fig. 1b). For the best-ancestral (BA) sets, we aimed at selecting the gene in each species with the highest sequence (BA-seq set) or expression (BA-expr set) conservation. For the best-divergent (BD) sets, we aimed at selecting the gene in each species with the lowest sequence (BD-seq set) or expression (BD-expr set) conservation. The results were overall highly consistent, in line with the elevated overlap between 1:1 sets **(Supplementary Fig. 1c)**, and with specific expected exceptions (see **Supplementary Discussion**).

### RNA-seq data and gene expression quantification

We used the bulk RNA-seq data from [10], which includes samples for the sixteen species in our phylogeny spanning up to eight homologous tissue types. We selected seven of those tissues (neural, testis, ovaries, muscle, excretory, epidermis and gut), excluding adipose because of the missing samples for the elephant shark. As an expression measure for each gene, we used the expression proportions per tissue (i.e., sum across tissues = 1). The expression proportion per tissue is defined as (tissue_expr / all_tissue_expr), where “tissue_expr” is the average quantile normalized log2(TPMs+1) expression of the gene in the target tissue and “all_tissue_expr” the sum of the average quantile normalized log2(TPMs+1) expression values across all tissues. See [10] for further details. All expression files are available in the **Supplementary Dataset**.

### Computation of sequence similarities

We derived two representative measures of sequence similarities for each gene orthogroup, one within vertebrates and one within insects. Each of these clade-level measures was computed following a 5-step procedure: (i) computation of pairwise protein sequence similarity between all possible pairs of orthologs, based on *BLOSUM62* and using pairwise protein alignments generated by *mafft* [51] with default parameters. Importantly, these pairwise protein sequence similarities were computed considering each gene first as query and then as target; (ii) for each query gene, computation of average sequence similarity between all its orthologs in each target species; (iii) for each query gene, computation of average sequence similarity among all target species (values from step (ii)); (iv) for each species, computation of the average sequence similarity between all its genes (values from step (iii)); (v) for each clade (vertebrates or insects), computation of the average sequence similarity between all its species (values from step (iv)). See **Supplementary Fig. 1a** for a detailed schematic of the procedure. Clade-level average sequence similarities for all gene orthogroups are provided in **Supplementary Table 1**. The sequence similarity values for the extra 1:1 orthogroup sets (represented in **Supplementary Fig. 4a,c,e,f, 5a,c,e,f, 6a,c,e,f, 7a,c,e,f**) were obtained with a similar procedure but skipping steps (i) and (ii) because of the presence of only one representative gene per species.

### Computation of expression similarities

We derived two representative measures of expression similarities for each gene orthogroup, one within vertebrates and one within insects. Each of these orthogroup-level measures was derived following a 5-step procedure: (i) computation of pairwise expression similarity between all possible pairs of orthologs. The pairwise expression similarity was defined as follows:

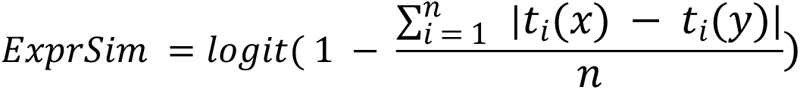

Where *n* represents the total number of considered tissue types (n=7), *i* indicates each of the tissues in turn, *x* and *y* correspond to the two species being compared. Briefly, we summed up the absolute differences in expression proportions across corresponding tissues, normalizing this value for the number of comparisons. As this measure would be inversely proportional to the conservation of expression profiles between two genes, we computed the actual expression similarity by subtracting it from one and applying a *logit* transformation. The resulting low and high expression similarity values correspond to gene pairs with greatly divergent and conserved expression profiles, respectively; (ii) for each query gene, computation of average expression similarity between all its orthologs in each target species; (iii) for each query gene, computation of average expression similarity among all target species (values from step (ii)); (iv) for each species, computation of the average expression similarity between all its genes (values from step (iii)); (v) for each clade (vertebrates or insects), computation of the average expression similarity between all its species (values from step (iv)). See **Supplementary** Fig. 1a for a detailed schematic of the procedure. Clade-level average expression similarities for all gene orthogroups are provided in **Supplementary Table 1**. The expression similarity values for the extra 1:1 orthogroup sets (represented in **Supplementary Fig. 4b,d,e,f, 5b,d,e,f, 6b,d,e,f, 7b,d,e,f**) were obtained with a similar procedure but skipping steps (i) and (ii) because of the presence of only one representative gene per species.

### Alternative measures for sequence and expression conservation

We first downloaded and formatted human phastCons scores (one file per chromosome) from UCSC: http://hgdownload.cse.ucsc.edu/goldenPath/hg38/phastCons100way/hg38.100way.phastCons/chr*.phastCons100way.wigFix.gz. PhastCons scores are nucleotide-level scores which represent the posterior probability of each nucleotide to be in its most-conserved state based on the comparison with 100 vertebrate genomes [52]. For each human gene belonging to our gene orthogroups, we computed the average phastCons values among all the nucleotides of its coding sequence. Finally, we compared these values with the average similarity values between the same query gene and all its vertebrate orthologs (see **Supplementary Fig. 3a**). Average phastCons value for each human gene are reported in **Supplementary Fig. 5**.

The expression correlations and expression distances (see **Supplementary Fig. 3b-l**) were based, respectively, on the Spearman’s correlations and the Euclidean distances (more precisely, 1 - Euclidean distance) of expression proportions across tissues. As for the sequence and expression similarities, we derived two representative measures of expression correlation/distance for each gene orthogroup, one within vertebrates and one within insects. These final measures were computed following the same 5-step procedure described for expression similarity (see above). See **Supplementary Fig. 1a** for a detailed schematic of the procedure. All average expression correlation/distance values are provided in **Supplementary Table 5**.

### Definition of housekeeping orthogroups

The definition of housekeeping genes was based on Tau, a measure of tissue-specificity ranging between 0 and 1 where low and high values correspond to broadly expressed and tissue-specific genes, respectively. Gene orthogroups were defined as housekeeping when all the conserved vertebrate genes presented a Tau value ≤ 0.25 (Tau values from [10]; 158 orthogroups in total, see **Supplementary Table 5**). Thus, in **Supplementary Fig. 3d-f** the expression similarities, correlations and distances are only shown for vertebrate orthogroups.

### Characterization of highly and lowly diversified gene orthogroups

For each clade (vertebrates and insects) and each considered feature (sequence and expression similarity) we defined one group of highly and one group of lowly diversified genes by selecting the 500 gene orthogroups with the lowest/highest similarity values (**Supplementary Table 2**). The phenotypes associated with highly diversified, lowly diversified and all orthogroups are defined as follows. First, we downloaded gene-phenotype associations from Ensembl ([53]; v105) for human and mouse and from FlyBase ([54] updated in January 2020) for the fruit fly. We then filtered only for the genes present in our gene orthogroups, and we created one vertebrate- and one insect-specific phenotypic annotation by associating the phenotype of the relative human/mouse or fly genes (respectively) to the whole orthogroup (see **Supplementary Dataset**). “Lethal” phenotypes were defined as everything containing “lethal” or “_die_”, “Not lethal” phenotypes corresponded to all other identified phenotypes, while all the orthogroups with no associated phenotype in the relative clade were labeled as NA. We then associated the phenotypic classification from the human/mouse and fly annotations to the vertebrates and insects gene groups, respectively, and plotted the resulting distributions (**Fig. 3b**). The number of species with duplications for each of these orthogroups (**Fig. 3c**) corresponded to the number of species in the relative clade with at least one paralog. The Tau values (**Fig. 3d**) were taken from [10]. The results depicted in **Supplementary Fig. 4g-i, 5g-i, 6g-i, 7g-i** were generated with the same procedure but starting from the additional 1:1 orthogroup sets.

### GO enrichments

The GO enrichments of common highly and lowly diversified gene groups (**Fig. 4b,c**) were performed using the human-transferred GO annotation from [10], and all gene orthogroups as a background. Common sets of highly and lowly diversified genes were defined as gene orthogroups included in at least three out of the four highly/lowly diversified groups (vertebrate/insect from the sequence/expression perspective). Up to the top 20 significant (FDR-corrected p-value ≤ 0.05) categories including at least 5 genes and 5% of the initial set are represented for each group in **Fig. 4a,b**, while all categories are reported in **Supplementary Table 3**. The GO enrichments for the extra 1:1 orthogroup sets (**Supplementary Fig. 4j, 5j, 6j, 7j**) were performed in the same way but using custom GO annotations that should better represent the gene selection in each particular orthogroup set. We generated these annotations by downloading the human GO annotation from Ensembl ([53]; v106) and transferring the function of the selected human gene to the relative orthogroup (similar to the approach described in [10] for the GO annotation used above).

### Gene Set Enrichment Analysis (GSEA)

We computed the deltas between the sequence and expression similarity measures in vertebrates and insects (**Supplementary Fig. 2a,b**), to which we applied a z-score transformation in order to shift the center of each distribution to zero (**Supplementary Fig. 2d,e**). We then used these z-scored deltas and the GO annotation previously described as input for two GSEAs, which were performed using the *fgsea* function in R (*fgsea* package) with the following parameters: *minSize=50, maxSize=500.* We then only filtered for the categories associated with an FDR-corrected p-value ≤ 0.01 and plotted the 30 categories with the highest/lowest normalized enrichment score (NES) in terms of either sequence or expression (**Fig. 5a**). All results are reported in **Supplementary Table 4)**. The genes highlighted in **Fig. 5b,c** are the ones listed under “leading edges” in the GSEA results. The GSEA depicted in **Supplementary Fig. 4k, 5k, 6k, 7k** were generated with the same procedure but starting from the different 1:1 orthogroup sets and using for each the custom GO annotation described above.

### Characterization of genes with diversification biases

In order to evaluate the over-representation of tissue-specific genes from particular tissues within the orthogroups with sequence or expression-driven specialization biases, we performed separate Fisher’s exact tests for each clade and each tissue. For each test, we ran the *fisher.test* function from base R with *alternative = “greater”* on a matrix containing (i) the number tissue-specific genes from the tested tissue included in the orthogroups with specialization biases, (ii) the total number of tissue-specific genes in the same orthogroups, (iii) the number of tissue-specific genes from the tested tissue in all other orthogroups and (iv) the total number of tissue-specific genes in all other orthogroups. The threshold for significant p-values was set at 0.05. The results depicted in **Supplementary** Fig. 4l, 5l, 6l, 7l were generated with the same procedure but starting from the different 1:1 orthogroup sets.

## Data availability

A Supplementary Dataset is available on Mendeley Data with DOI: 10.17632/vdxfrvvb3w.1.

## Code availability

The code used for all analyses and figure generation is provided in a Github repository at https://github.com/fedemantica/vertebrate_insect_GE.

## Authors contribution

FM and MI conceived the study. FM generated all results and figures. FM and MI wrote the manuscript.

## Acknowledgements

The tissue icons appearing in Fig.1a are original drawings by Queralt Tolosa Ramon, subject to a CC BY-NC-SA (Attribution-NonCommercial-ShareAlike) 4.0 International license. Animal silhouettes were generated with *Bing Chat* by MIcrosoft (2023) https://www.bing.com/search.

## Supplementary Discussion

In this Supplementary Discussion, we aim to support the robustness of our study with two different approaches. First, we will compare our selected measures of sequence and expression similarity with alternative conservation metrics, describing the specific advantages associated with our approach. Second, while we believe that considering all genes in each orthogroup was the best way to capture its overall variability, we will argue that selection of representative orthologs does not affect our main conclusions.

### Comparison between measures of sequence conservation

The estimation of sequence conservation is at the core of several comparative studies. A common measure to estimate such conservation between groups of orthologous sequences is phastCons, a nucleotide-level conservation score [55,56] from UCSC [57]. However, as effective as it is in many other contexts, this score was optimal for our dataset. First, phastCons scores are derived from a multiple alignment of a query genome to a set of target genomes. These scores are available in UCSC for only 5 out of 8 of our vertebrate species, and they were all derived from the comparison with different sets of target vertebrate genomes. Moreover, phastCons metrics were released for only one of the insect species in our dataset (*Drosophila melanogaster*), and they were mainly determined by the comparison with other Drosophila genomes. Thus, the use of these particular measures would likely introduce a species-related bias to our results, as (i) not all the species would be equally represented and (ii) the available scores were derived from comparison with different sets of target genomes.

Given these limitations, we implemented a simple measure of sequence similarity between orthologous genes based on protein alignments, adjusted to our dataset and biological questions. This similarity measure presents two main advantages: first, all species in a given orthogroup have the same weight in determining the final similarity values, removing eventual species-specific biases; second, these values reflect the overall level of molecular diversification in a given gene orthogroup (not only the variation eventually under evolutionary selection) and are directly comparable across gene families. Moreover, we indeed detected a good correlation between this sequence similarity measure and the relative average phastCons score at the level of single human genes (see **Supplementary Fig. 3a** and **Methods**), suggesting that our value of sequence similarity is largely concordant with more standard metrics for the species in which these are available.

### Comparison between measures of expression conservation

Previous works investigating the evolution of sequence and expression divergence in closely related animal groups have returned somewhat contradicting results depending on the adopted measure of expression divergence. In some cases, a correlation between the two features was detected [28,31,32]. In other cases, this association did not emerge [33,34], likely due to the issues linked to the use of correlations as a metrics for expression divergence [35].

In order to test how our results are affected by the use of distinct conservation measures, we also computed expression correlations and expression distances (based on Spearman’s correlations and Euclidean distances, respectively; see **Supplementary Fig. 3b,c**, **Methods** and **Supplementary Table 5**). As for the expression similarities, both these metrics showed global overlapping distributions between vertebrates and insects (**Supplementary Fig. 3b,c**). However, specific analyses of house-keeping genes suggest that, unlike expression similarities and distances, expression correlations do not accurately reflect the conservation level of the underlying expression profiles (**Supplementary Fig. 3d-f**). Moreover, while the expression distances captured all the patterns identified by the expression similarities (**Supplementary Fig. 3g-i**), none of them were retrieved by the use of expression correlations (**Supplementary Fig. 3j-l**).

In conclusion, we find that both expression similarity and distance serve as more effective measures of expression divergences compared to expression correlations, at least when a limited number of conditions are being compared.

### Comparison between different representative 1:1 orthogroup sets

In order to test if and how the selection of representative genes in each species affects the key results in our study, obtained considering the values for all paralogs within an orthogroup and species, we generated four distinct 1:1 gene orthogroup sets, each composed of paralogs picked according to different conservation criteria. In particular, the best-ancestral (BA) orthogroup sets contain the gene in each species characterized by the highest sequence (BA-seq set) or expression (BA-expr) conservation; on the contrary, the best-divergent (BD) orthogroup sets comprised the most divergent gene per species in terms of either sequence (BD-seq set) or expression (BD-expr set) (see **Methods** and **Supplementary Fig. 1b**). Importantly, these orthogroup sets are highly overlapping, as around 75% of the representative genes in each of them are also contained in all the others (**Supplementary Fig. 1c**).

In general, the distributions of sequence and expression similarities across orthogroups and clades for all 1:1 orthogroup sets are highly overlapping to the original ones using all paralogs (see **Fig.1d,e** and **Supplementary Fig. 4a,b, 5a,b, 6a,b, 7a,b**), as well as their inter-clade and intra-clade correlations (see **Fig.2** and **Supplementary Fig. 4c-f, 5c-f, 6c-f, 7c-f**). However, we detected some subtle differences between 1:1 orthogroup sets, which were in line with the nature of the selected representative genes. In fact, sequence similarity of the BA-seq orthogroups showed higher correlation between vertebrates and insects (**Supplementary Fig. 4c**) compared to when whole orthogroups are used (**Fig. 2a**). The same is true for the expression similarity of the BA-expr orthogroups (**Supplementary Fig. 6c** vs **Fig. 2b**), while the opposite pattern is observed for the BD-seq (**Supplementary Fig. 5a**) and BD-expr (**Supplementary Fig. 7c**) orthogroups. Thus, even if the general patterns we described by using all orthogroups are robust to ortholog selection, this procedure introduced some small changes that are in line with the nature of the orthogroups themselves (i.e., higher concordance between clades and conservation measures when the most conserved orthologs are selected).

In addition, the results relative to the molecular features associated with higher diversification levels (**Fig. 3b-d**; i.e., higher tissue-specificity, increased gene duplication and lower representation of essential genes) were highly consistent across all 1:1 orthogroup sets (**Supplementary Fig. 4g-i, 5g-i, 6g-i, 7g-i**). The same is true for the enrichment in house-keeping functions and molecular machinery for the common lowly diversified gene orthogroups. However, we obtained slightly different results for the common highly diversified best-divergent orthogroups, which either did not show any significant enrichment (BD-seq) or showed only a couple of significant categories related to extracellular space (BD-expr) (**Supplementary Fig. 4j, 5j, 6j, 7j**). These different results were again consistent with the nature of the selected orthologs, as the most divergent genes of the BD sets are more likely to reflect random molecular variation compared to the most conserved orthologs or to when the whole orthogroup is considered.

Finally, we performed a GSEA (same as in **Fig. 5a**) to characterize the functional profile of the gene orthogroups with clade-specific diversification biases (**Supplementary Fig. 4k,l, 5k,l, 6k,l, 7k,l**). In all cases, insect genes with a diversification bias showed significant enrichments for testis-specific genes belonging to cilium-related categories, consistent with the results reported for the whole orthogroups. Also genes with diversification bias in vertebrates revealed consistent enrichment for neural-related genes and functional categories, even if these enrichments were not as strong in the BA-expr set. This was partially expected, as this particular set was by definition the one less likely to capture tissue-specific genes.

In conclusion, the results presented in this study are highly robust to ortholog selection procedure, and the general patterns we uncovered reflect true intrinsic properties of a given gene family that are captured independently from which gene(s) the analyses are based on.

## Supplementary figures

**Supplementary Fig. 1:**
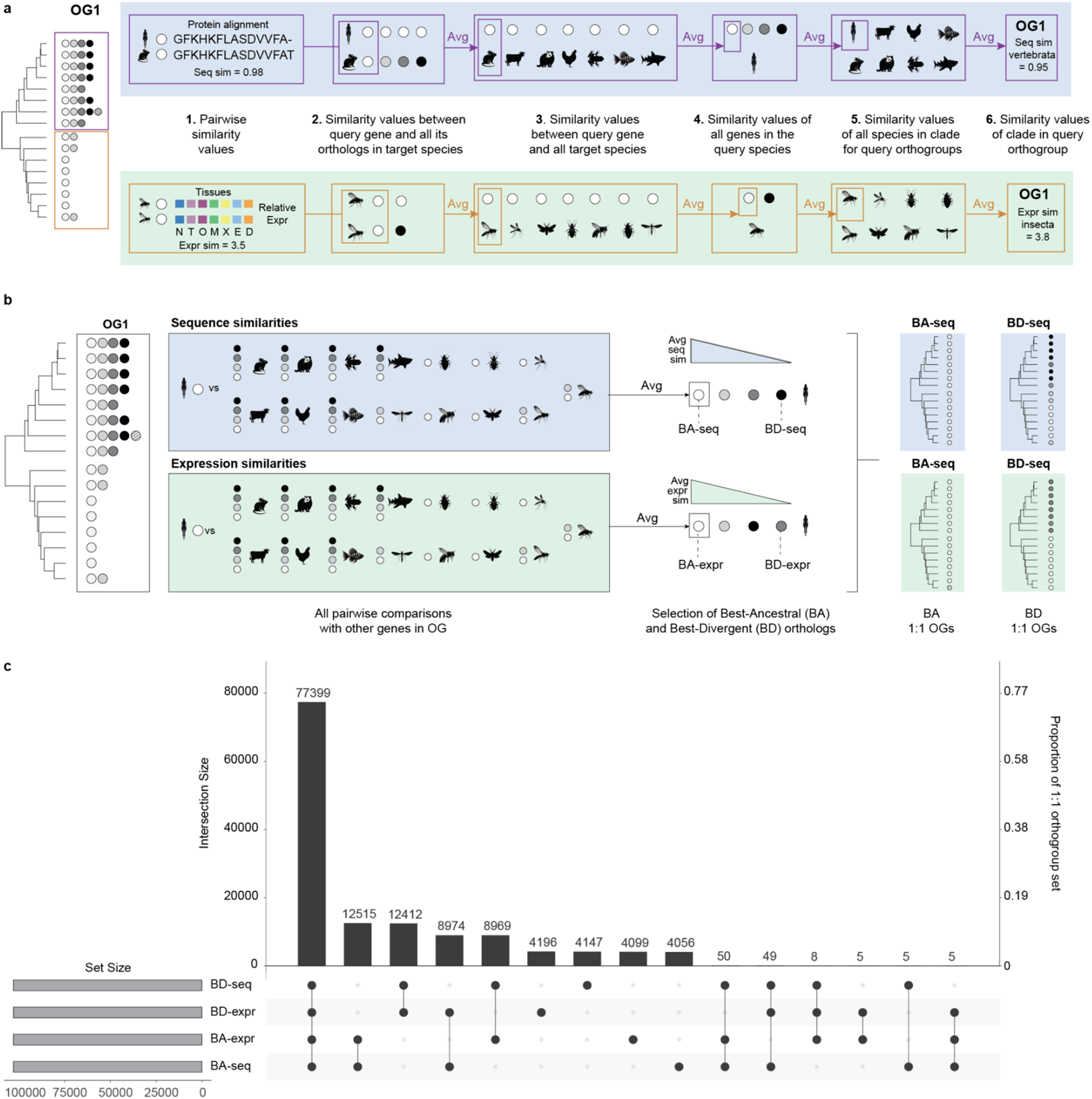
Schematics for computation of sequence/expression similarity and definition of extra 1:1 orthogroup sets. **a.** Example schematic for the computation of sequence similarities (top) or expression similarities (bottom) for each orthogroup and each clade. **b.** Scheme for the definition of all extra 1:1 orthogroup sets (BA-seq, BA-expr, BD-seq, BD-expr. see Methods). **c.** UpSet plot showing the overlap among all extra 1:1 orthogroup sets. **Abbreviations:** Avg, Average; OGs, orthogroups.

**Supplementary Fig. 2:**
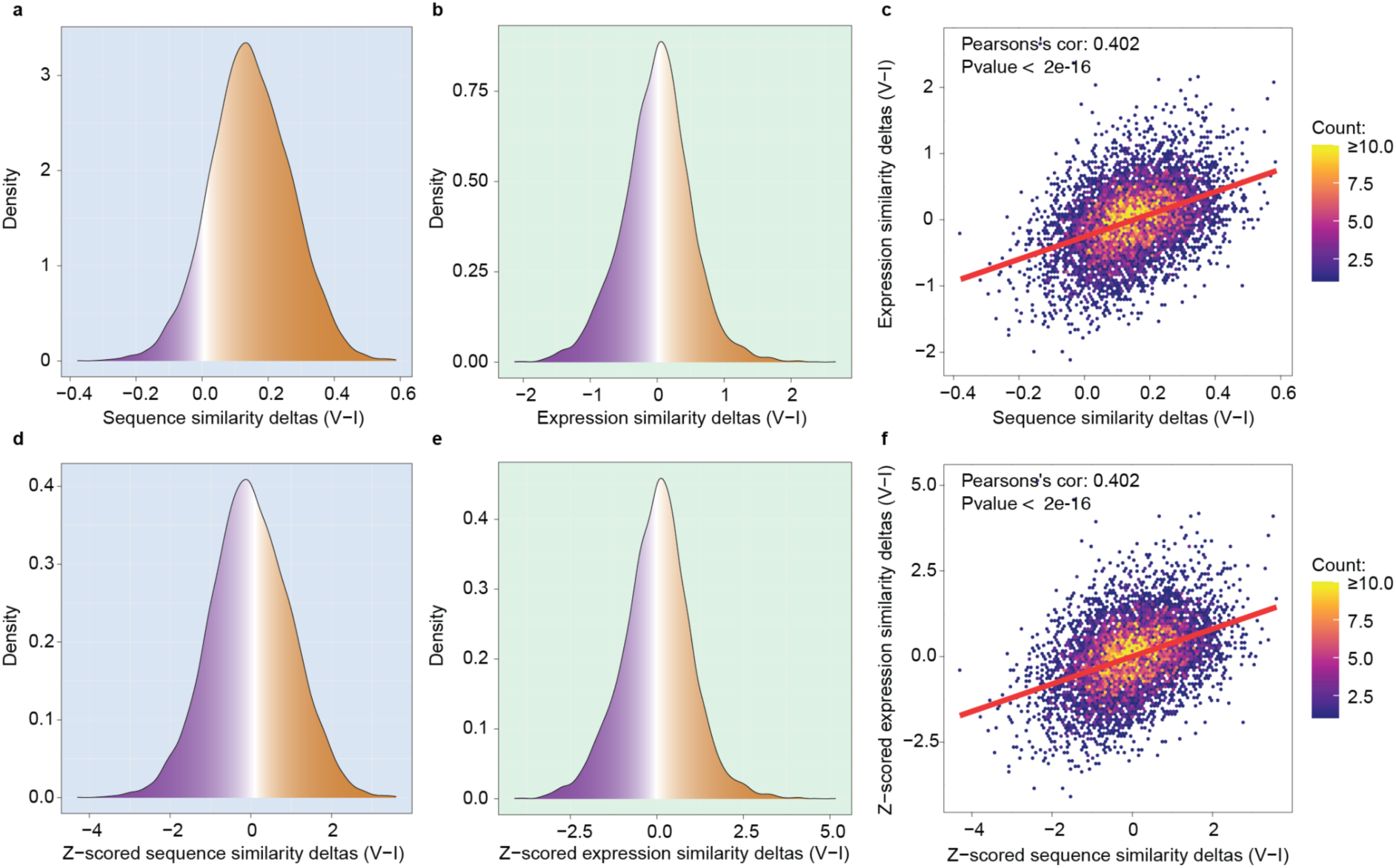
Deltas of sequence and expression similarities between vertebrates and insects. **a,b.** Distribution of sequence similarity deltas (a) and expression similarity deltas (b) between vertebrates and insects for all gene orthogroups (n=6,787). Color gradients indicate orthogroups with negative (purple) or positive (orange) delta values. **c.** Correlation of the sequence similarity deltas (x axis) and expression similarity deltas (y axis) across all gene orthgroups (n=6,787). **d,e.** Distribution of z-scored sequence similarity deltas (d) and z-scored expression similarity deltas (e) between vertebrates and insects for all gene orthogroups (n=6,787). Color gradients indicate orthogroups with negative (purple) or positive (orange) delta values. **f.** Correlation of the z-scored sequence similarity deltas (x axis) and z-scored expression similarity deltas (y axis) across all gene orthgroups (n=6,787).

**Supplementary Fig. 3:**
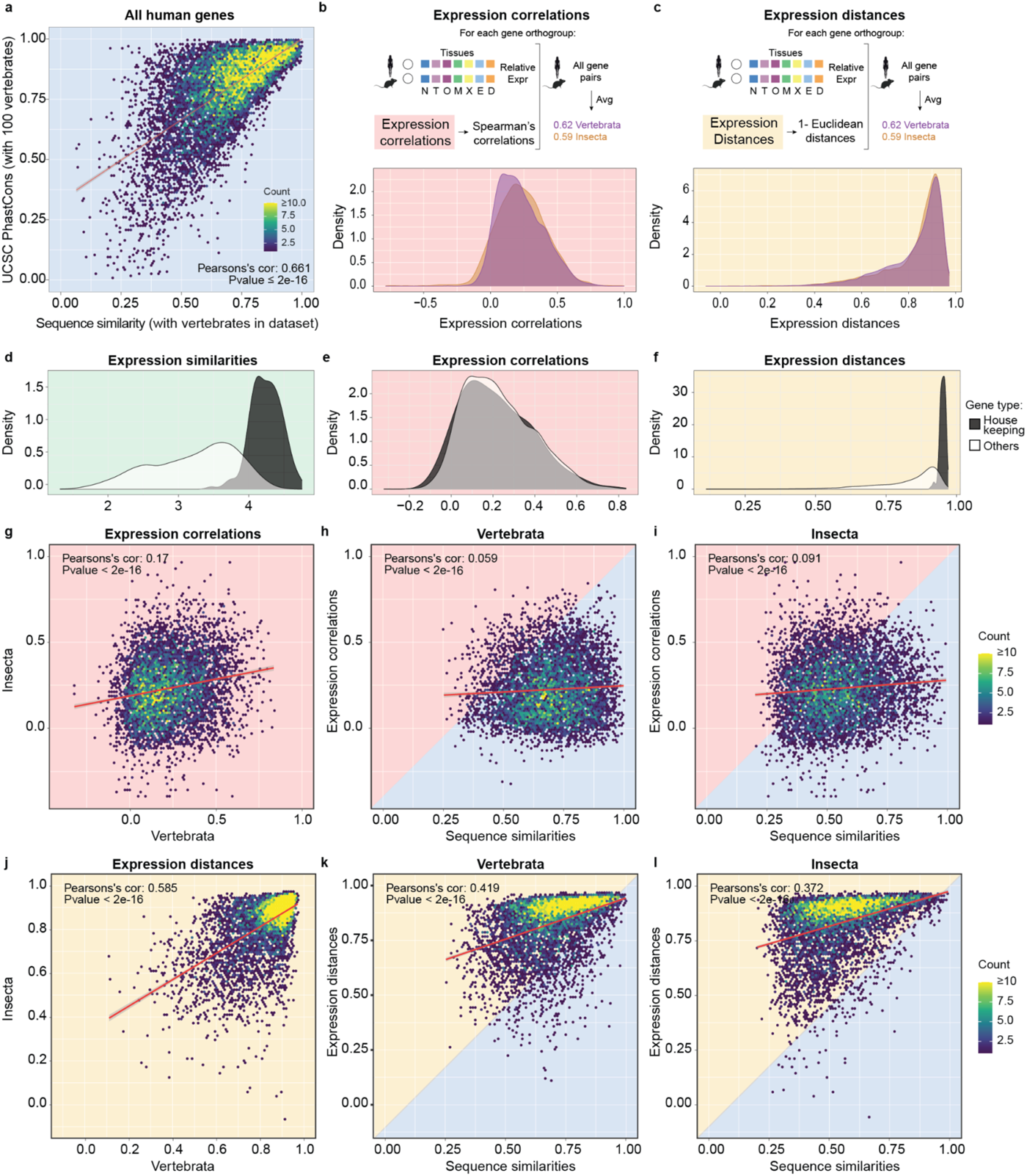
Comparison between alternative measures of sequence and expression conservation. **a.** Correlation of average sequence similarity (x axis) of each human gene with its average phastCons score (y axis; see **Methods** for definition of average phastCons). **b,c.** top: scheme for the computation of expression correlations (b) and expression distances (c). See **Supplementary** Fig. 1a for step-by-step schematics (same logic as the expression similarity). Bottom: distributions of the expression correlations (b) and distances (c) values for all gene orthogroups (n=6,787), with vertebrate and insect measures plotted in purple and orange, respectively. **d,e,f.** Distribution of expression similarities (d), correlations (e) and distances (f) within vertebrates divided between housekeeping (black) and other genes (white). See **Methods** for definition of housekeeping genes. **g.** Correlation of the expression correlations of all gene orthogroups between vertebrates (x axis) and insects (y axis) (n=6,787). **h,i.** Correlation between the sequence similarities (x axis) and expression correlations (y axis) of all gene orthogroups within vertebrates (h) and insects (i) (n=6,787). **j.** Correlation of the expression distances of all gene orthogroups between vertebrates (x axis) and insects (y axis) (n=6,787). **k,l.** Correlation between the sequence similarities (x axis) and expression distances (y axis) of all gene orthogroups within vertebrates (k) and insects (l) (n=6,787).

**Supplementary Fig. 4:**
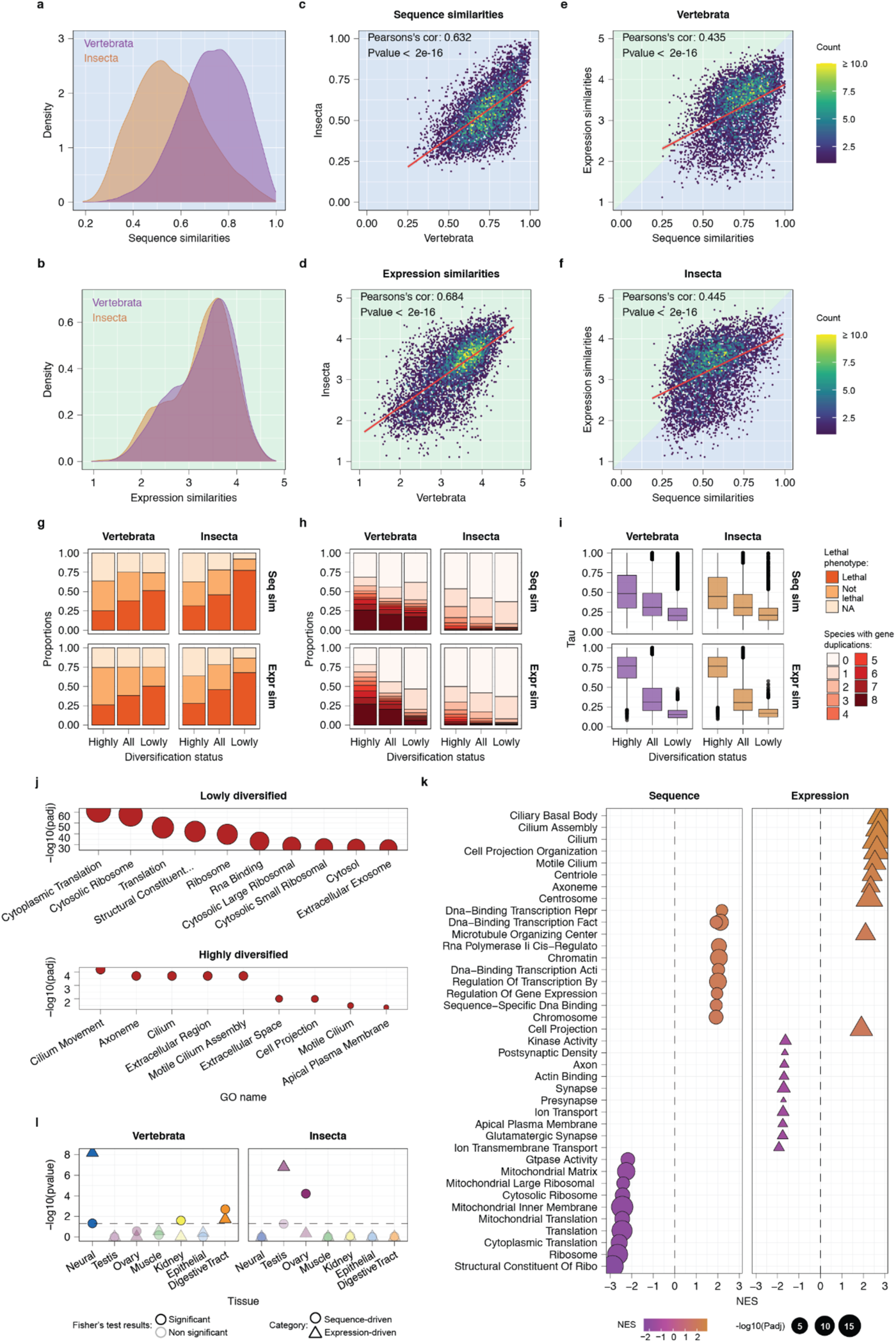
Key results repeated with the BA-seq 1:1 orthogroup set. **a, b.** Distributions of the sequence (a) and expression (b) similarity values for all gene orthogroups (n=6,787) within vertebrates (purple) and insects (orange). **c,d.** Correlation of the sequence (c) and expression (d) similarities of all gene orthogroups between vertebrates (x axis) and insects (y axis) (n=6,787). **e, f.** Correlation between the sequence (x axis) and expression (y axis) similarities of all gene orthogroups within vertebrates (e) and insects (f) (n=6,787). **g.** Proportions of highly diversified, all and lowly diversified orthogroups associated with a lethal, non-lethal and uncharacterized phenotype. Legend on the side of panel i. **h.** Duplication profiles of highly diversified, all, and lowly diversified orthogroups, where species with duplications are defined as all species with at least two conserved paralogs in a given orthogroup. Legend on the side of panel i. **i.** Distribution of Tau values for genes included in highly diversified, all and lowly diversified gene orthogroups. **j.** Significant categories (FDR corrected p-values ≤ 0.05, intersection ≥ 5, precision ≥ 0.05) from a GO enrichment analysis performed on the lowly (top) and highly (bottom) diversified gene defined as in Fig. 4b,d. Results are limited to the top 10 categories. **k.** Significant categories (FDR-corrected p-values ≤ 0.01) from a GSEA performed on the z-scored deltas of sequence (left) or expression (right) similarities between vertebrates and insects. Results are limited to the top 10 categories with highest and lowest NES for each feature (sequence and expression) and clade (vertebrates and insects) **l.** Results of Fisher’s exact tests from the *fisher.test* function in R (*alternative=”greater”*) comparing the proportions of tissue-specific genes with diversification biases (Tau ≥ 0.75) in each tissue to the same proportions in the background. Tests with p-values ≤ 0.05 are considered as significant. Abbreviations: NES, normalized enrichment score.

**Supplementary Fig. 5:**
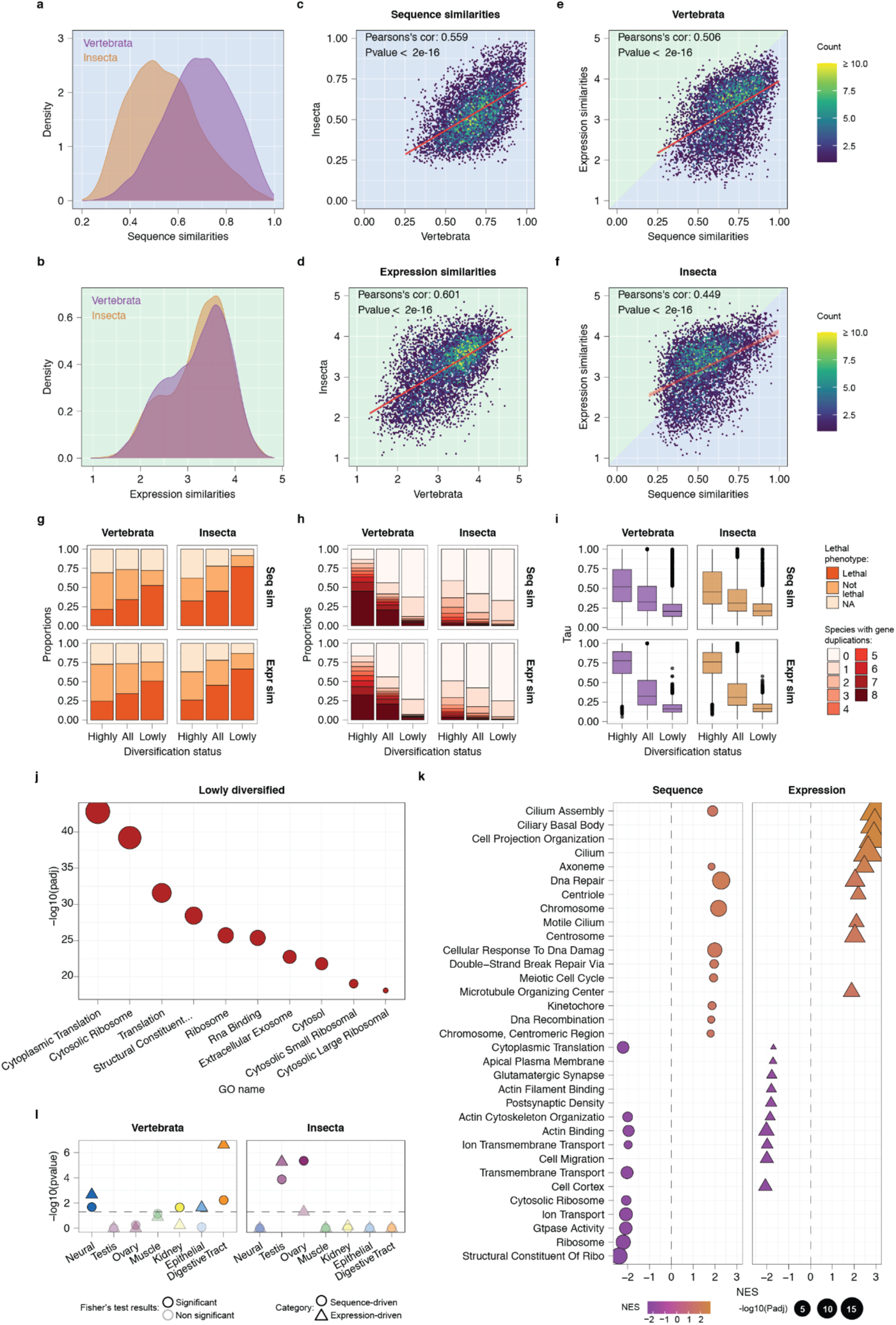
Key results repeated with the BD-seq 1:1 orthogroup set. **a, b.** Distributions of the sequence (a) and expression (b) similarity values for all gene orthogroups (n=6,787) within vertebrates (purple) and insects (orange). **c,d.** Correlation of the sequence (c) and expression (d) similarities of all gene orthogroups between vertebrates (x axis) and insects (y axis) (n=6,787). **e, f.** Correlation between the sequence (x axis) and expression (y axis) similarities of all gene orthogroups within vertebrates (e) and insects (f) (n=6,787). **g.** Proportions of highly diversified, all and lowly diversified orthogroups associated with a lethal, non-lethal and uncharacterized phenotype. Legend on the side of panel i. **h.** Duplication profiles of highly diversified, all, and lowly diversified orthogroups, where species with duplications are defined as all species with at least two conserved paralogs in a given orthogroup. Legend on the side of panel i. **i.** Distribution of Tau values for genes included in highly diversified, all and lowly diversified gene orthogroups. **j.** Significant categories (FDR corrected p-values ≤ 0.05, intersection ≥ 5, precision ≥ 0.05) from a GO enrichment analysis performed on the lowly (top) and highly (bottom) diversified gene defined as in Fig. 4b,d. Results are limited to the top 10 categories. **k.** Significant categories (FDR-corrected p-values ≤ 0.01) from a GSEA performed on the z-scored deltas of sequence (left) or expression (right) similarities between vertebrates and insects. Results are limited to the top 10 categories with highest and lowest NES for each feature (sequence and expression) and clade (vertebrates and insects) **l.** Results of Fisher’s exact tests from the *fisher.test* function in R (*alternative=”greater”*) comparing the proportions of tissue-specific genes with diversification biases (Tau ≥ 0.75) in each tissue to the same proportions in the background. Tests with p-values ≤ 0.05 are considered as significant. Abbreviations: NES, normalized enrichment score.

**Supplementary Fig. 6:**
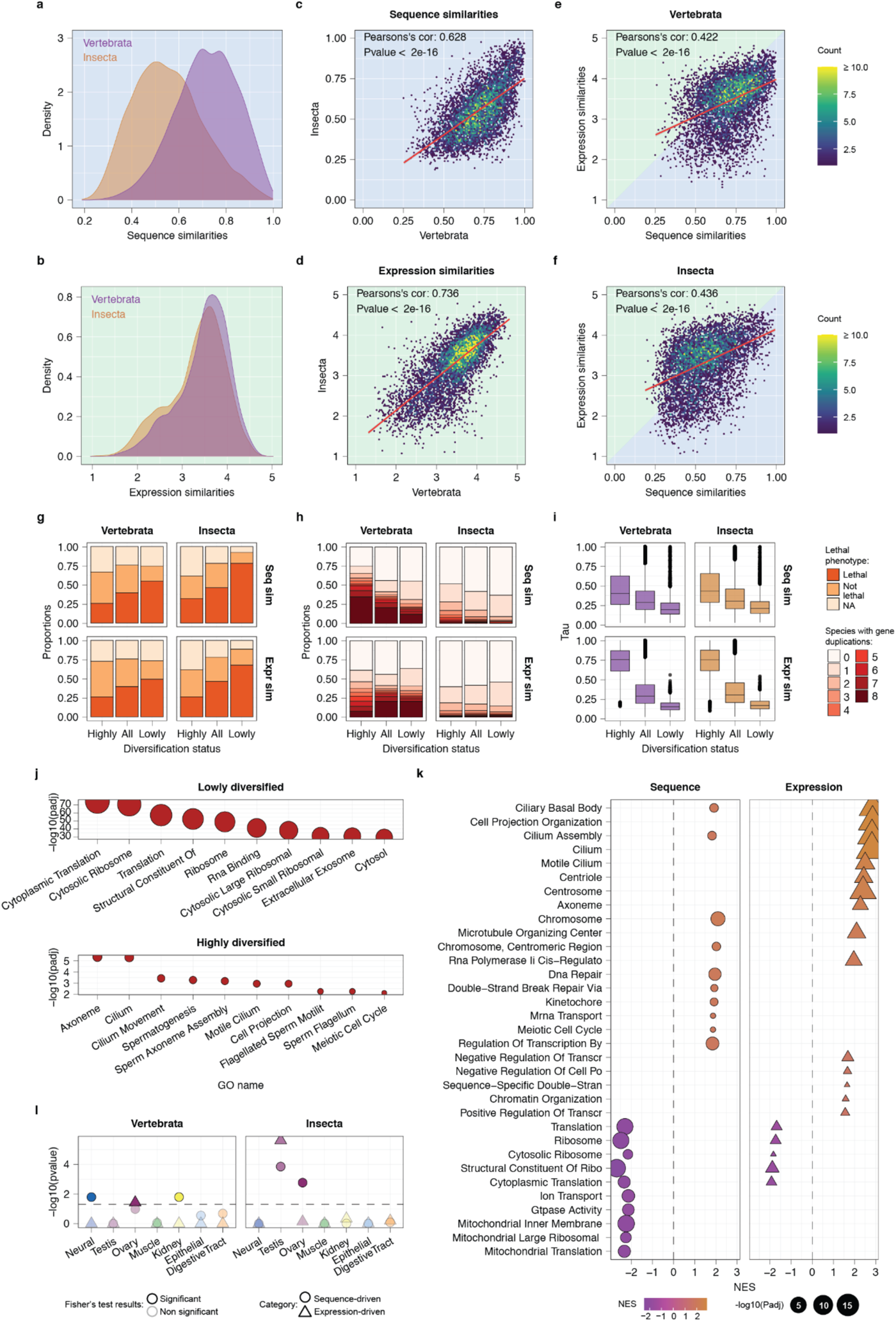
Key results repeated with the BA-expr 1:1 orthogroup set. **a, b.** Distributions of the sequence (a) and expression (b) similarity values for all gene orthogroups (n=6,787) within vertebrates (purple) and insects (orange). **c,d.** Correlation of the sequence (c) and expression (d) similarities of all gene orthogroups between vertebrates (x axis) and insects (y axis) (n=6,787). **e, f.** Correlation between the sequence (x axis) and expression (y axis) similarities of all gene orthogroups within vertebrates (e) and insects (f) (n=6,787). **g.** Proportions of highly diversified, all and lowly diversified orthogroups associated with a lethal, non-lethal and uncharacterized phenotype. Legend on the side of panel i. **h.** Duplication profiles of highly diversified, all, and lowly diversified orthogroups, where species with duplications are defined as all species with at least two conserved paralogs in a given orthogroup. Legend on the side of panel i. **i.** Distribution of Tau values for genes included in highly diversified, all and lowly diversified gene orthogroups. **j.** Significant categories (FDR corrected p-values ≤ 0.05, intersection ≥ 5, precision ≥ 0.05) from a GO enrichment analysis performed on the lowly (top) and highly (bottom) diversified gene defined as in Fig. 4b,d. Results are limited to the top 10 categories. **k.** Significant categories (FDR-corrected p-values ≤ 0.01) from a GSEA performed on the z-scored deltas of sequence (left) or expression (right) similarities between vertebrates and insects. Results are limited to the top 10 categories with highest and lowest NES for each feature (sequence and expression) and clade (vertebrates and insects) **l.** Results of Fisher’s exact tests from the *fisher.test* function in R (*alternative=”greater”*) comparing the proportions of tissue-specific genes with diversification biases (Tau ≥ 0.75) in each tissue to the same proportions in the background. Tests with p-values ≤ 0.05 are considered as significant. Abbreviations: NES, normalized enrichment score.

**Supplementary Fig. 7:**
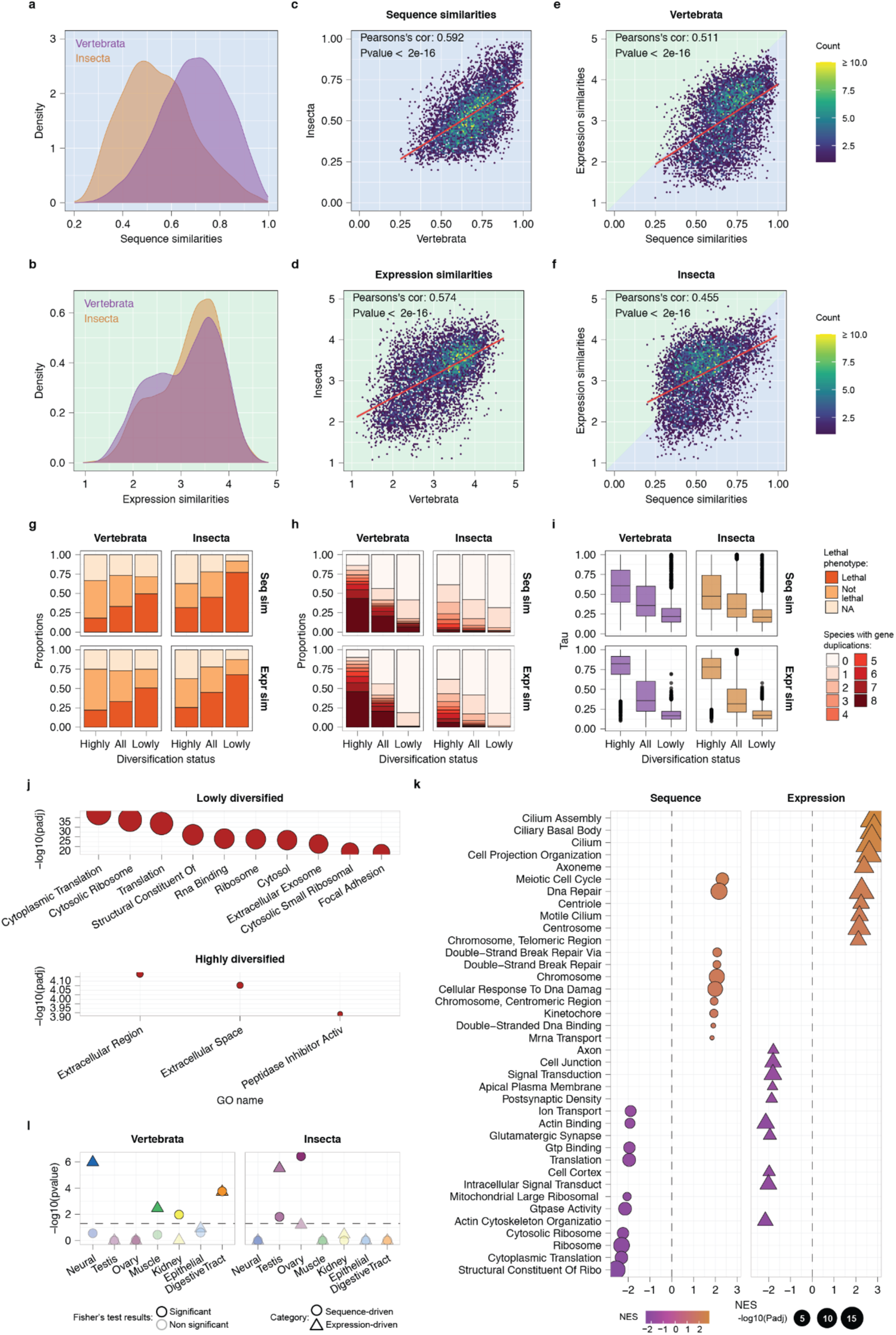
Key results repeated with the BD-expr 1:1 orthogroup set. **a, b.** Distributions of the sequence (a) and expression (b) similarity values for all gene orthogroups (n=6,787) within vertebrates (purple) and insects (orange). **c,d.** Correlation of the sequence (c) and expression (d) similarities of all gene orthogroups between vertebrates (x axis) and insects (y axis) (n=6,787). **e, f.** Correlation between the sequence (x axis) and expression (y axis) similarities of all gene orthogroups within vertebrates (e) and insects (f) (n=6,787). **g.** Proportions of highly diversified, all and lowly diversified orthogroups associated with a lethal, non-lethal and uncharacterized phenotype. Legend on the side of panel i. **h.** Duplication profiles of highly diversified, all, and lowly diversified orthogroups, where species with duplications are defined as all species with at least two conserved paralogs in a given orthogroup. Legend on the side of panel i. **i.** Distribution of Tau values for genes included in highly diversified, all and lowly diversified gene orthogroups. **j.** Significant categories (FDR corrected p-values ≤ 0.05, intersection ≥ 5, precision ≥ 0.05) from a GO enrichment analysis performed on the lowly (top) and highly (bottom) diversified gene defined as in Fig. 4b,d. Results are limited to the top 10 categories. **k.** Significant categories (FDR-corrected p-values ≤ 0.01) from a GSEA performed on the z-scored deltas of sequence (left) or expression (right) similarities between vertebrates and insects. Results are limited to the top 10 categories with highest and lowest NES for each feature (sequence and expression) and clade (vertebrates and insects) **l.** Results of Fisher’s exact tests from the *fisher.test* function in R (*alternative=”greater”*) comparing the proportions of tissue-specific genes with diversification biases (Tau ≥ 0.75) in each tissue to the same proportions in the background. Tests with p-values ≤ 0.05 are considered as significant. Abbreviations: NES, normalized enrichment score.

